# Every cloud has a silver lining: Moderate plant water stress positively impacts development, immunity, and gut microbiota of a specialist herbivore

**DOI:** 10.1101/411678

**Authors:** E Rosa, G Minard, J Lindholm, M Saastamoinen

## Abstract

The ongoing global temperature rise has led to increasing frequency of drought events, negatively impacting vegetation and the living organisms relying on it. Extreme drought killing host plants can clearly reduce herbivore fitness, but the impact of moderate host plant water stress on insect herbivores can vary, and may even be beneficial. The Finnish Glanville fritillary butterfly (*Melitaea cinxia*) has faced reduced precipitation in recent years, which has impacted population dynamics. However, whether the negative effects depend on extreme desiccation killing the host plant or moderate drought impacting plant quality remains unclear. We assessed the performance of larvae fed on moderately water-stressed *Plantago lanceolata* in terms of growth, gut microbial composition and immune response. We found that larvae fed on water-stressed plants had better growth, a more heterogeneous bacterial community and a shifted fungal community in the gut, and up-regulated the expression of one candidate immune gene (pelle), whereas survival remained unaffected. Most of the measured traits showed considerable variation due to family structure. Our data suggest that in temperate regions moderate host plant water stress can positively shape resource acquisition of this specialized insect herbivore, potentially by increasing nutrient accessibility or concentration. Potentially, the better larval performance may be mediated by a shift of the microbiota on water-stressed plants, calling for further research especially on the understudied gut fungal community.

## Introduction

The present-day escalating greenhouse gas emissions have led to global climate change [1]. One manifestation of this change is the increasing occurrence of events of drought or low precipitation [1,2], impairing the biomass and species richness of plants, and consequently of the herbivore species relying on them [3,4]. Insects can be differently affected by plant drought, depending on the duration and frequency of the stress, and on the insect feeding guild or species [5]. However, continuous water stress is generally expected to have a negative impact [5]. Nonetheless, the degree of the water stress is also important, and extreme desiccation killing the host plant is likely to have a negative impact on herbivore fitness. Importantly, insects should not be considered individually, but rather as holobionts composed of the insect host and its microbial community [6]. Microbial communities play a role in insect digestion, immunity and development, amongst others [7–11]. Microbiota composition varies with host habitat and food source [12]. Contrarily to other insects, the impact and spread of intracellular symbionts is quite limited in Lepidoptera [13], with recent work suggesting that bacterial communities are often acquired through food and environment [13,14]. Therefore, changes in the plant microbiota due to climatic factors might lead to drastic changes in the microbiota of larvae.

Despite a large body of literature on the effects of plant condition on insect development, the impact on other indicators of insect condition, as the ability to maintain an adequate immune response, and a balanced gut microbial composition remain largely unexplored. Insect immunity can vary in relation to diet abundance and quality [15,16], presence of plant defense compounds [17–19], and microbial composition of the plant [reviewed in 20]. Moreover, energetic costs due to the activation of insect immunity are expected to rise on suboptimal diets [21].

We used the Finnish Glanville fritillary butterfly as model system to investigate the effects of water-stressed host plants on the performance of larvae feeding on them. Based on estimates from the field and long-term data [22,23], reduced precipitation is expected to substantially impact population dynamics. Pre-diapause larvae are very sessile and have limited ability to move among host plants. Hence, their survival strongly depends on the condition of the host plant they develop on. Whether the suggested poorer performance of larvae during drought events is due to starvation due to extreme plant desiccation, or also to subtler changes in host plant quality driven by moderate water stress remains unknown. We reared a set of pre-diapause larvae on host plants subject to a moderate continuous water stress or constantly well-watered, and compared their performance. We assessed larval performance in terms of growth rate, immunity, survival, and diversity of the gut microbial community (bacteria and fungi).

## Material and methods

### Experimental design

The seeds of the host plant, *Plantago lanceolata*, were collected in the Åland Islands in 2015. The plants were reared in a greenhouse with artificial lightning (16/8h L/D). We introduced a moderate water stress on the host plants as follows: a batch of 74 plants was daily watered with 20 ml for the “water stress” treatment, whereas a batch of 38 plants was watered with 40 ml for the “well-watered” treatment. We used *M. cinxia* larvae of eleven separate families, which were the F1 of individuals collected in Åland in 2015. Forty 24h-old larvae per family were assigned to each treatment (water stress vs. well-watered), and reared in sterile petri dishes. Larvae were fed daily a 2.25 cm^2^ leaf piece corresponding to their given treatment until the 4^th^ instar, while 4.5 cm^2^ leaf pieces were used after the 4^th^ instar. To prevent short-term dryness, 0.35 ± 0.05 ml of water was added on the surface of the leaf pieces. Developmental time was estimated as the date when 1/2 of the larvae within a petri dish molted to a new instar. Survival was assessed once larvae reached the 5^th^ and diapausing instar. At least 25 larvae per treatment per family survived until the 5^th^ and diapausing instar, except for one family. At this stage, all larvae were weighed, and five randomly chosen were snap-frozen and stored at -80°C for immune gene expression. To introduce an immune challenge for the phenoloxidase assay, we punctured with a microneedle the penultimate proleg of 20 diapausing larvae after 24h of starvation: ten were challenged with *Micrococcus luteus* (250 mg/ml lyophilized cells), and ten with PBS (Phosphate-buffer saline solution, Gibco-USA) as control. Twenty-four hours later these larvae were snap-frozen and stored at -80°C for further analysis.

### Microbial community structure

All larvae challenged with *M. luteus* and PBS were individually washed three times with 500µl of 1X PBS (Ambion-USA), surface-sterilized in 70% ethanol and washed five times in 1X PBS. Midguts were dissected under sterile conditions in 1X PBS. Ten pooled midguts were used per condition per family. DNA was extracted from pooled midguts using Qiagen DNeasy Blood and Tissue kit (Qiagen-Germany) with a previously adapted protocol [24]. Bacterial and fungal Automated Ribosomal Intergenic Spacer Analysis (ARISA) were performed with primers amplifying the 16S-23S rDNA (bacteria) and 18S-28S rDNA (fungi) intergenic sequences. Bacterial ARISA (b-ARISA) was carried out in triplicates for each sample with the primers ITSF (5’FAM -GTC GTA ACA AGG TAG CCG TA-3’), ITSReub (5’-GCC AAG GCA TCC ACC-3’) amplifying the 16S-23S rDNA intergenic sequence and the fungal ARISA was conducted with the primers 2234C (5’HEX-GTT CCG TAG GTG AAC CTG C-3’) and 3126T (5’-ATA TGC TTA AGT TCA GCG GGT-3’) amplifying the 18S-28S rDNA intergenic sequence [25,26]. PCR was conducted with a reaction mixture containing 1X of Q5 buffer (New England Biolabs, USA), 1X of High-GC enhancer (New England Biolabs, USA), 200 µM of dNTP (Thermo Fisher scientific, USA), 500 nM of each primer and a total of 0.12 mg×ml^-1^ of Bovine Serum Albumin (New England Biolabs, USA), 0.06 mg×ml^-1^ of T4gene32 (New England Biolabs, USA) in a final volume of 25 µl. Amplifications were conducted on an ABI Veriti thermal cycler (Applied Biosystem, USA) with 3min of denaturation at 94 °C, 30 cycles of 45 s at 95 °C, 1 min at 55 °C, 1 min 20 s at 72°C followed by a final elongation of 1 min 20 s at 72°C. Replicates of each sample were first controlled on a 1% agarose gel for positive amplification, then purified with a PCR Clean-up kit (Macherey-Nagel, Germany) following the manufacturers’ recommendations, quantified with NanoDrop (ThermoFisher) and diluted at a concentration of 10 ng×µl^-1^. Capillary migrations were performed on a 3730XL Bioanalyzer (Applied Biosystem, USA), using 4 µl of each sample, 10.8 µl of Hi-Di formamide and 0.2 µl of GS 1,200 LIZ size marker (Thermo Fisher scientific, USA). The fluorograms were analyzed with the software Genemapper 4.0. The signals comprised between 100-1000 bp were selected, binned to windows of 5 bp and transformed into relative fluorescence intensities (RFI) using a previously published method [27].

### Phenoloxidase activity

The phenoloxidase enzyme is an indicator of insect immunocompetence [28]. As gut dissection caused complete bleeding, we could not restrict the assay to the haemocoelic PO activation alone, but we assayed whole body homogenate of ten pooled larval carcasses. The measurements were based on modified methods by [28]. Pools of ten larval carcasses per family per treatment were weighed and placed in a screw-cap tube with 20 µl of ice-cold PBS per mg measured and crushed with a metal bead in a Bead-ruptor machine (30 rps for 1.5 min). The tubes were centrifuged at 13 000 rpm for 10 min at 4°C, and 20 µl of the supernatant were transferred in a new Eppendorf tube with 20 µl of ice-cold PBS. The assay was performed in 96-well plates and the increase in absorbance due to melanin production was measured every minute for 100 min at 490 nm and 30 °C with an EnSpire microplate reader (PerkinElmer). Each well included 70 µl of ice-cold distilled water, 10 µl of ice-cold PBS, 7.5 µl of sample, and 10 µl of 10 mM L-Dopa solution. The PO activity was assessed as the slope of the reaction curve during the linear phase (V_max_).

### Immune gene expression

Five individuals per family per plant watering condition were assessed for immune gene expression. Typically, immune responses are tested with an immune challenge serving as positive control. Due to data limitation, this was not possible; hence immune gene expression is assessed only in response to plant water stress. However, previous work shows upregulation of most of the tested genes under bacterial challenge [29]. The qPCR was performed on individual carcasses dissected as described above. RNA was extracted by homogenization of the samples in 1ml of Trisure (Bioline-Germany; Appendix 1). The qPCR targeted seven immune genes following methods by [29]: Lysozyme C, prophenoloxidase, attacin, peptidoglycan recognition protein LC, β-1,3-glucan recognition protein, serpin 3a and pelle, as well as two housekeeping genes: mitochondrial ribosomal protein L37 and S24. Phase separation was performed by supplementing the mixture with 200 µl of Chloroform and centrifuging the mixture at 12 000 g for 15 min at 4°C. The aqueous phase was then collected, precipitated in 0.5 ml of isopropyl alcohol for 10 min at 12 000g and 4°C for 10 min. The RNA pellet was washed with 1 ml of 75% ethanol and centrifuged for 5 min at 4°C. RNA pellets were dried and dissolved in 20 µl of DEPC-treated water (Ambion, USA). Dissolved RNAs were then treated with DNAse I (Thermo Fisher Scientific, USA) and reverse transcribed into cDNA using iScript cDNA Synthesis Kit (Bio-rad, USA) and the manufacturer’s conditions.

The qRT-PCR values were transformed into a relative expression according to the 2^-δδCt^ method using the housekeeping genes Ct values and the immunity genes Ct values of individuals fed on well-watered plants as baseline.

### Statistical analysis

The b-ARISA and f-ARISA normalized matrix were used as an input to compare the bacterial and fungal communities among the midguts of *M. cinxia.* The diversity analysis were conducted with the package *vegan* in R [30,31]. The α-diversity comparisons were conducted with the Shannon index using linear mixed models and the family was used as a random effect. The β-diversity analyses were conducted using the Bray-Curtis distance. Unconstrained ordinations of the Bray-Curtis community distances among the samples was represented using a non-metric multidimensional scaling (NMDS) and the potential correlation between waterstress plant treatment, phenoloxidase activity and the community structure was tested on the individuals with a permutational analysis of variance and plotted with a Canonical Analysis of Principal Coordinates (CAP). The effect of the plant treatment on growth, survival, phenoloxidase activity and immune gene expression were estimated with a Linear Mixed Model approach with treatment as explanatory variable and family as a random variable. We tested the impact of the random effect ‘family’ by calculating intraclass correlation coefficients (ICC) of all the models as proxy of the proportion of phenotypic variation explained by larval genetic background. All models were run with packages *ade4* and *LmerTest* in R [31–33].

## Results

Larvae had a markedly higher growth rate on water-stressed compared to well-watered host plants (*F*_1,29_=9.5, *P*=0.004; Fig 1), while survival was unaffected by diet (*P*>0.3). The variance explained by larval family was ∼20% for growth rate and over 30% for survival (Table 1). The α-diversity of fungal and bacterial communities associated with *M. cinxia* did not respond to bacterial challenge or water stress treatment (Table 1). However, the fungal community was shifted (*adonis*-ANOVA; F_1,36_=1.425; *P*=0.026; Fig 2) and the bacterial community was more heterogeneous (HOMOVA; F_1,38_=10.071; *P*=0.008; Fig. 3) in larvae fed with water-stressed plants, with no effect of bacterial challenge (Table 1). Larval family considerably impacted α-diversity (39.32% and 52.90% for bacterial and fungal communities, respectively), but only moderately affected its composition (8.37% and 25.70% for bacterial and fungal communities, respectively).

**Table 1.**
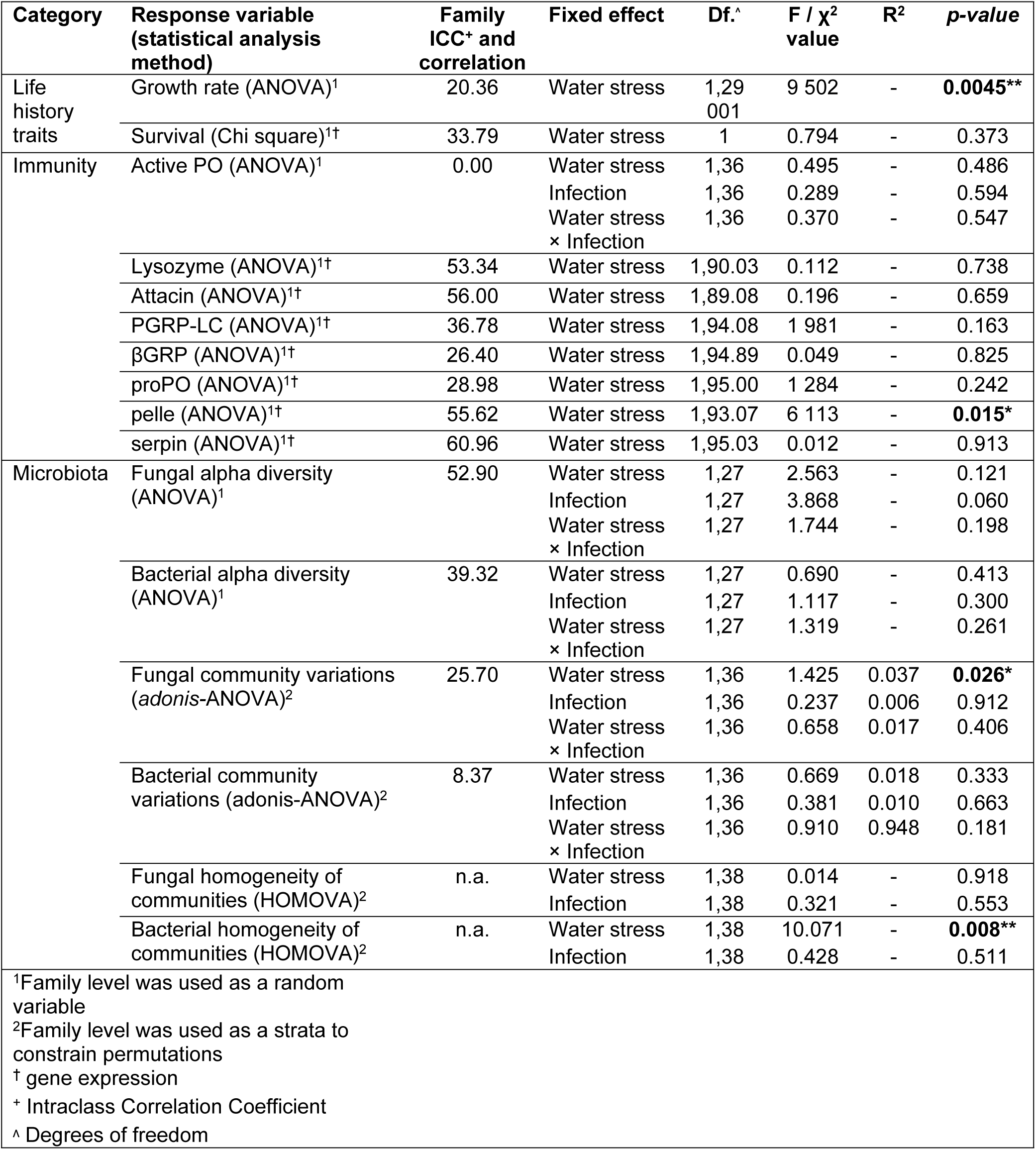
Effect of plant water stress and bacterial infection on the larval life-history traits, immunity and microbiota of *M. cinxia*

**Figure 1.**
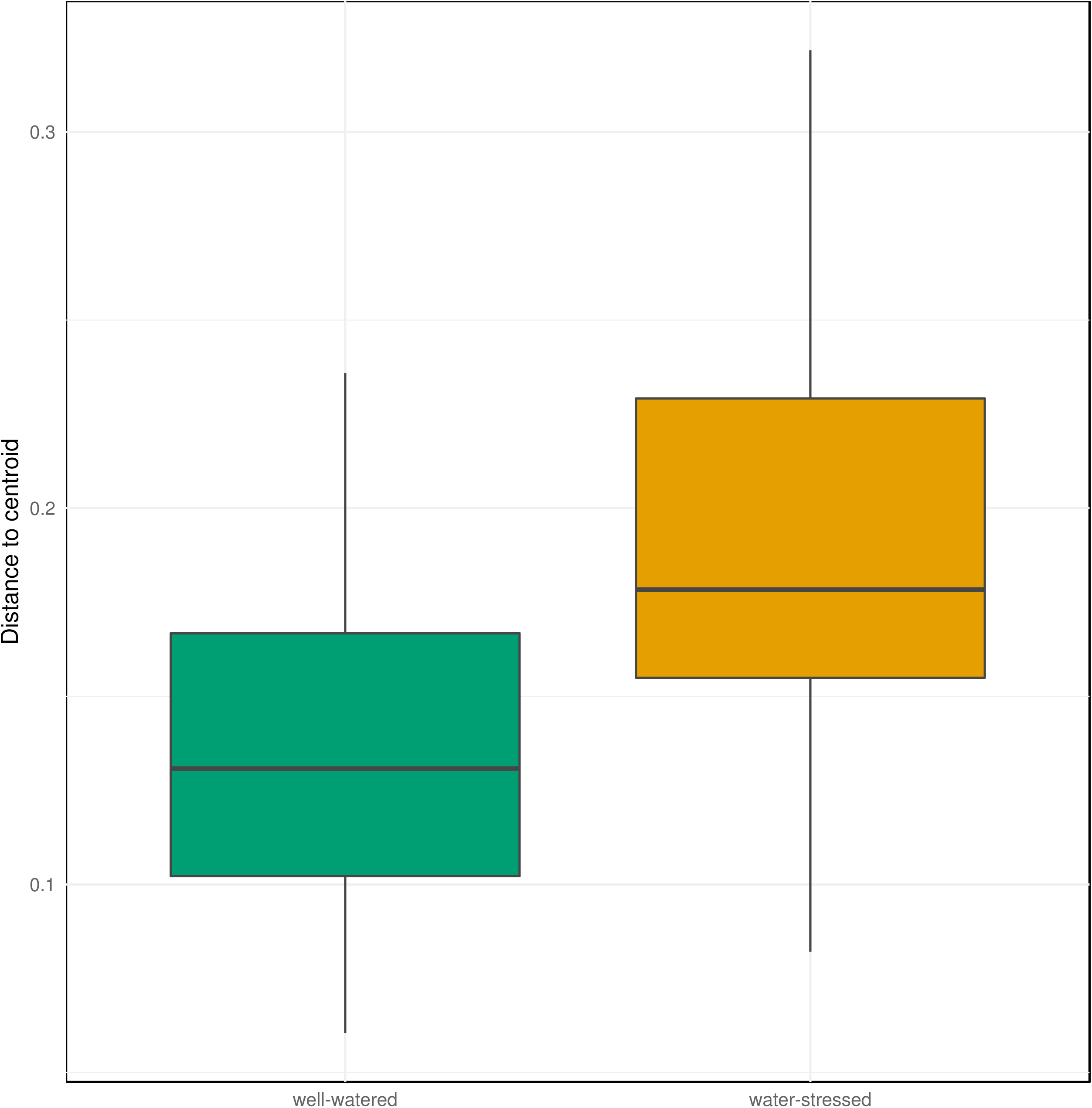
Larval growth rate of *Melitaea cinxia* on well-watered and water-stressed host plants. Larvae fed on well-watered and water-stressed host plant are displayed in green and orange fill, respectively.

**Figure 2.**
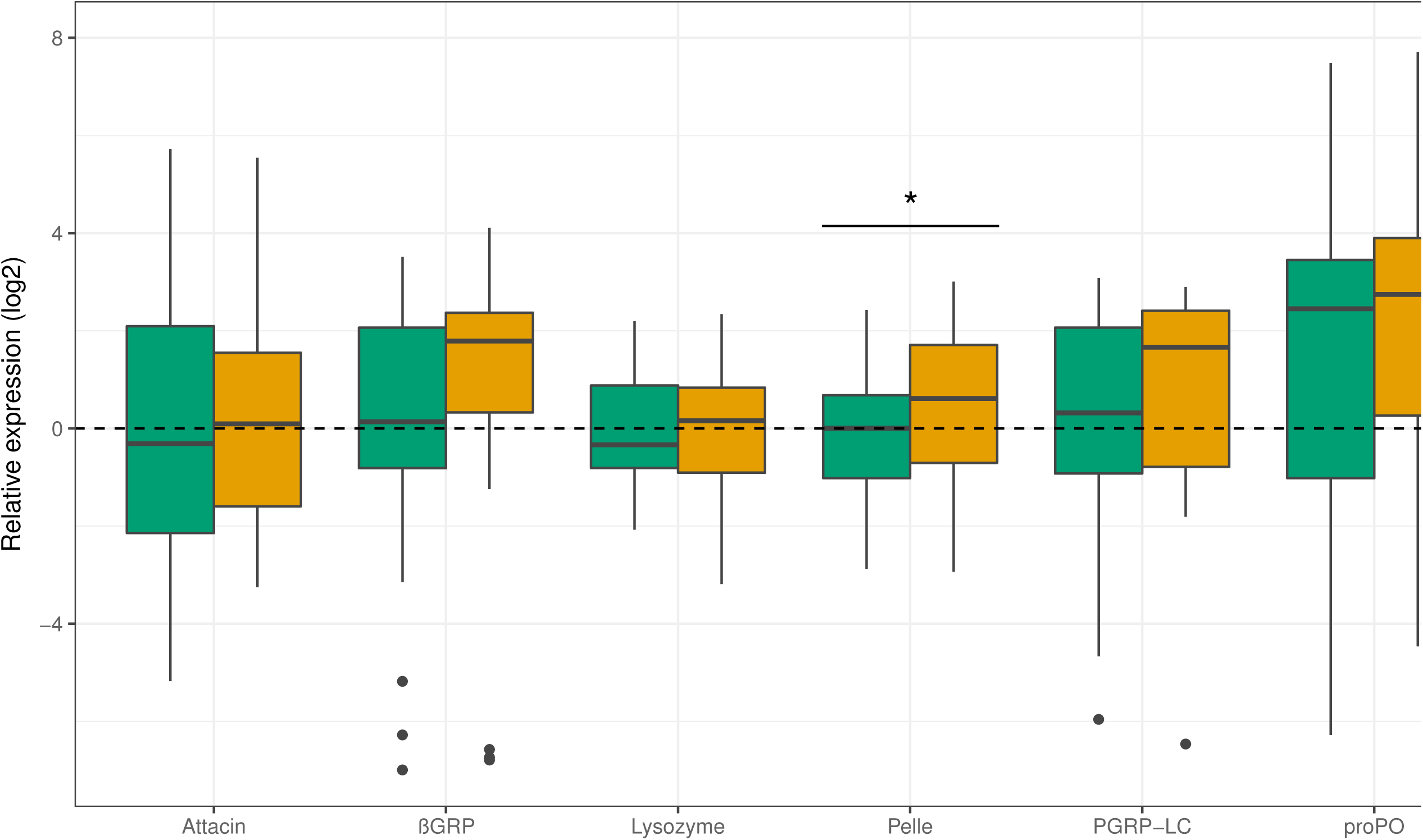
Canonical Analysis of Principal coordinates (CAP) of the fungal β-diversity among guts of *Melitaea cinxia* fed with well-watered or water-stressed host plants. Larvae fed on well-watered or water-stressed host plants are represented in green and orange respectively.

**Figure 3.**
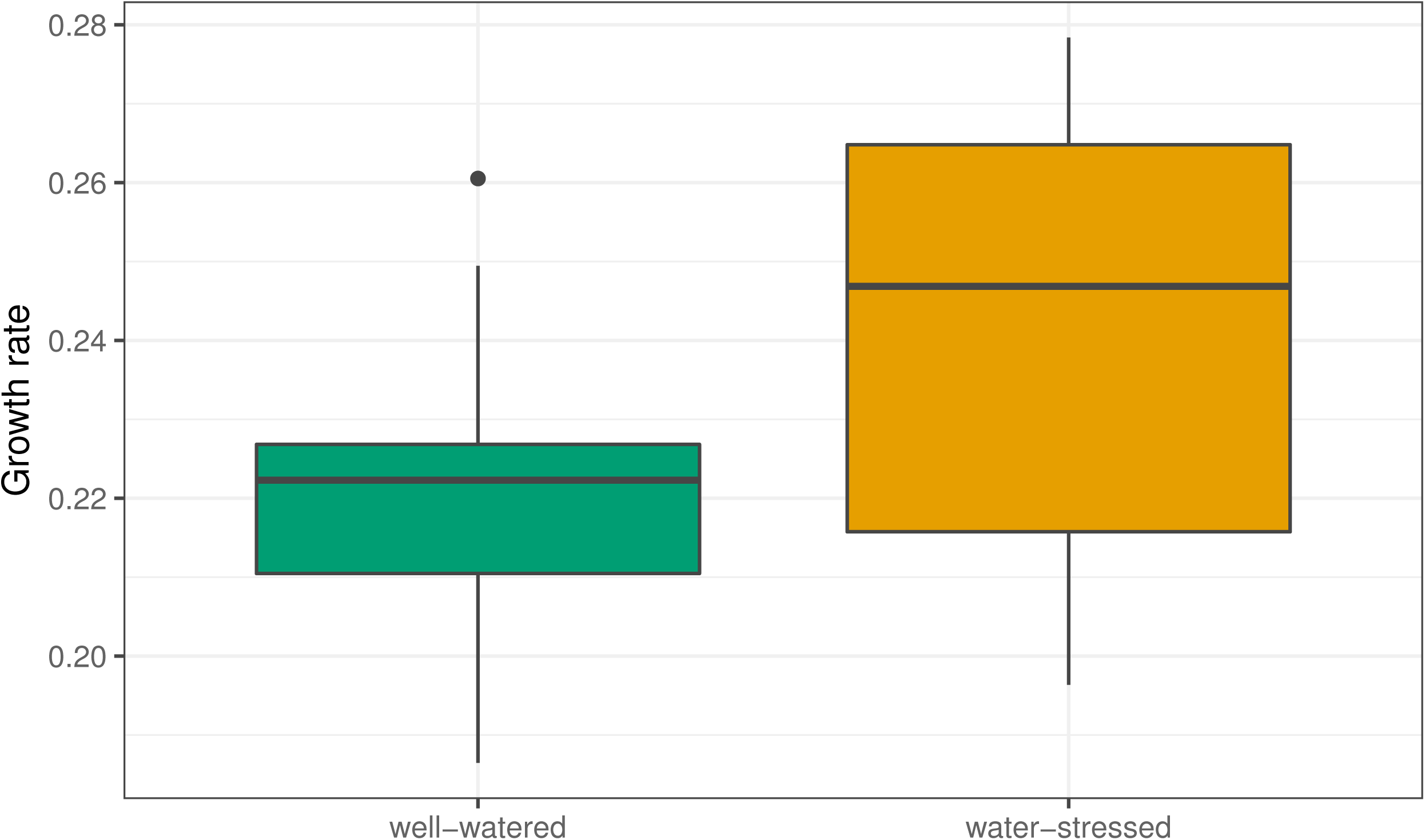
Heterogeneity of the bacterial community in the gut of *Melitaea cinxia*. Distances from the group centroid represent the heterogeneity of the bacterial community within each groups. Larvae fed on well-watered or water-stressed host plants are represented in green and orange respectively.

PO activity was unaffected by larval diet or bacterial challenge (*P*<0.4 for both, Table 1). Pelle expression was upregulated by host plant water stress (*F*_1,93_=6.1, *P*<0.015; Table 1; Fig 4), while the remaining genes were unaffected (*P*>0.1 for all). Larval family considerably affected the variance in immune gene expression (Table 1).

**Figure 4.**
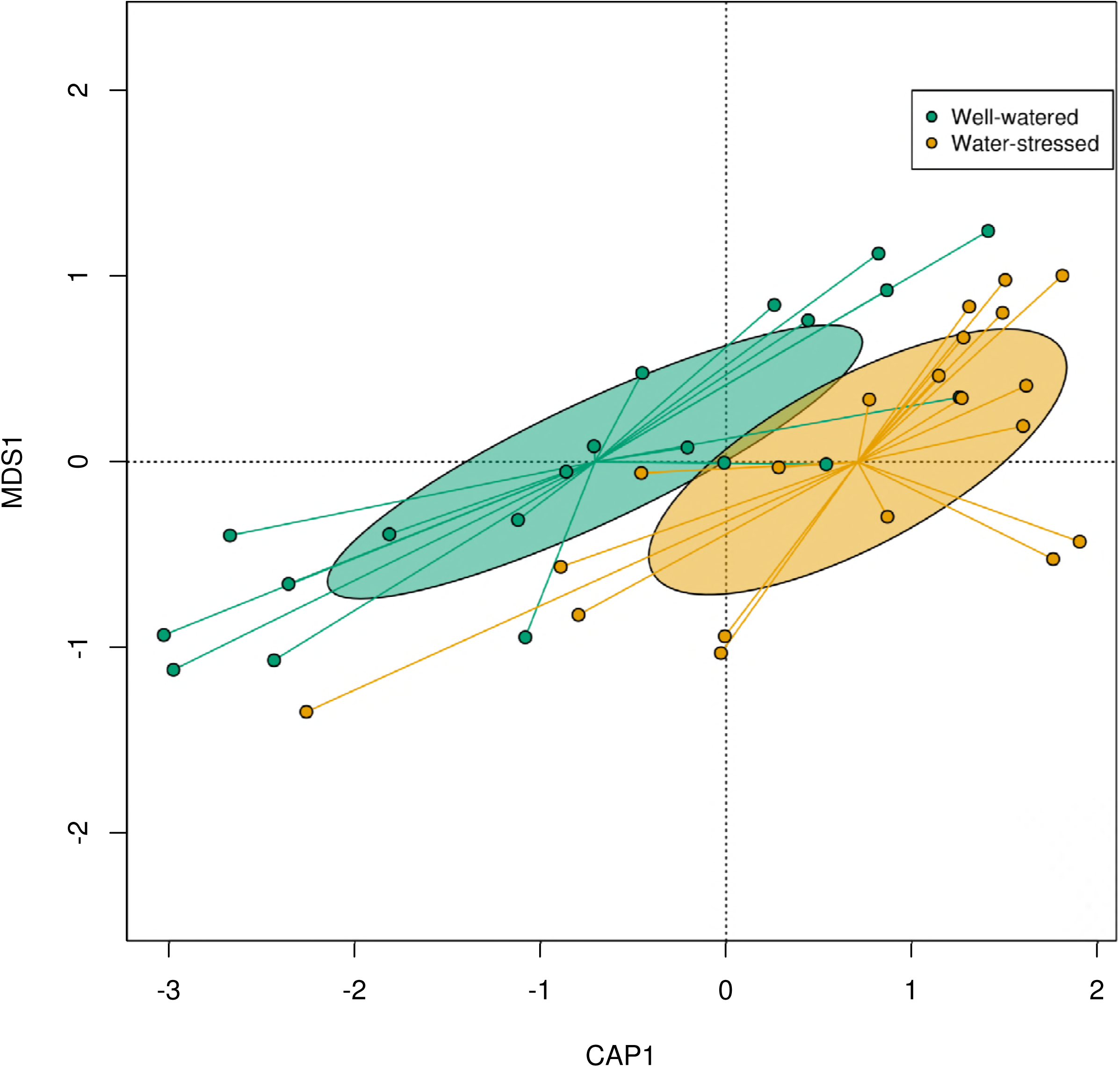
Immune gene expression of larvae fed on well-watered and water-stressed host plants. Larvae fed on well-watered and water-stressed host plants are displayed in green and orange fill, respectively.

## Discussion

In line with the present day global warming trends, recent work has shown that population dynamics of the Glanville fritillary butterfly can be heavily impacted by drought or low precipitation events [22], highlighting the relevance of water stress in shaping natural communities. We found that larvae reared on mildly water-stressed host plants grew better with no impact on survival. The microbial community of guts was more diverse in larvae fed on water-stressed plants, and this may have played a role in positively shaping larval growth. One of the immune genes assessed, coding for pelle, was upregulated in larvae fed with water-stressed plants. We are unaware of the implications of this response, but the activation of an immune reaction toward infective microbes ingested with the food seems unlikely, given the absence of fitness costs on the water stress diet. In general, our findings suggest a higher larval performance on moderately water-stressed plants. Intriguingly, larvae of the same age but faced with severe plant water stress have a low performance in terms of both reduced survival and body mass [34].

One well-documented effect of water stress on plants is an increase in the accumulation of amino acids and sugars in leaf tissues [35,36]. Potentially, the better growth rate of larvae found here could be explained in terms of increased nutrient accumulation or accessibility on water-stressed plants. However, this should be confirmed by future work formally quantifying macronutrient content in leaf tissues. Interestingly, larvae fed on water-stressed and well-watered plants had some differences in their gut microbiota composition: the bacterial community was more heterogeneous in the water stress treatment, while the fungal community was shifted, meaning that the fungal taxa composing the gut community differed almost completely between the two feeding treatments. In most Lepidoptera, the bacterial gut microbiota is mostly acquired through the food and is hence mostly transient [14]. Therefore, gut bacteria are suggested to have little impact on individual performance [14]. Yet, in few cases bacterial symbionts have been shown to impact insect reproduction or their ability to be protected against pathogens [13]. Conversely, eukaryotic microorganisms interacting with Lepidoptera such as fungi are currently poorly characterized, and their impact on host performance is largely unknown. However, studies with other insect models suggest that fungi might play a role in nutrient-provisioning [37], steroid synthesis [38] or protection against pathogens [39]. Intestinal microorganisms are also known to modulate gut immunity of insect hosts [40]. In turn, insects control their microbiota composition by modulating specific immune pathways [40]. The effect of plant drought on fungal microbes found here suggests exciting perspective for future research on insect-plant interactions. With the present data, we are unable to say, however, whether the impact on microbiota composition and immune gene expression is driven directly by the food ingested, or indirectly via some microbiota-immunity interaction.

The immune gene coding for pelle was upregulated by larvae feeding on water-stressed host plants. Pelle is a protein of the Toll pathway, which is responsive to fungi, yeast and gram-positive bacteria [41]. Pelle is involved in the release of antimicrobial peptides (AMPs), defensive molecules which are only expressed upon infection [41]. We are unaware whether the activation of pelle eventually led to the production of antimicrobial peptides as the only candidate AMP gene tested here, attacin, is effective against gram-negative bacteria [42], and hence not part of the Toll pathway featuring pelle. Indeed, the mechanism linking plant water stress and AMPs production remains to be explored, and its implications can only be speculated. On one hand, this may indicate that some potentially harmful microbes were detected in larvae fed on water-stressed plants, activating pelle and AMP production. The microbes causing this response may have been ingested with the food or their relative abundance in the gut may have been altered by the ingestion of water-stressed plant tissues. However, larvae fed on water-stressed plants had higher performance, which is unlikely in the presence of infectious microbes. On the other hand, larvae may have been only primed by microbes found on the water stress diet, but without the activation of an immune reaction featuring AMP production. In order to understand the significance of this response, future work involving detailed characterization of the microbial community taxa of both diets is needed.

To conclude, we found mostly beneficial effects of moderate plant drought-stress on insect performance, indicating that the previously suggested negative impact of reduced precipitation is likely due to extreme desiccation [22,34]. Indeed, this pattern is confirmed by work finding reduced survival and body mass of larvae fed on severely water-stressed plants [34]. Our water stress treatment may in fact reflect a rather common condition in the field, where the butterfly often occupies open and dry outcrop meadows [43]. Notably, we found a more heterogeneous and diverse gut microbial community under the water stress treatment, and we suggest it potentially plaid a role in shaping the positive larval performance observed in terms of growth rate. Finally, the shift in the fugal community in response to diet detected here calls for future research on the understudied eukaryotic microbial community interacting with lepidopteran hosts.

## Acknowledgements

We thank Suvi Ikonen for helping during the experiment, Aapo Kahilainen for designing the water stress treatment and Toshka Nyman for performing the qPCR analysis.

## Ethical statement

Insects and plants are not legally concerned by ethical regulations.

## Data accessibility

Data can be found in Dryad (doi:10.5061/dryad.86b77t3)

https://datadryad.org/review?doi=doi:10.5061/dryad.86b77t3

## Authors’ contributions

ER, GM, JL and MS designed the experiment. JL conducted the experiment. ER, GM and JL analyzed the data. All authors equally contributed to the final version of the manuscript.

